# Equivalent Change Enrichment Analysis: Assessing Equivalent and Inverse Change in Biological Pathways between Diverse Experiments

**DOI:** 10.1101/586875

**Authors:** Jeffrey A. Thompson, Devin C. Koestler

## Abstract

*In silico* functional genomics have become a driving force in the way we interpret and use gene expression data, enabling researchers to understand which biological pathways are likely to be affected by the treatments or conditions being studied. There are many approaches to functional genomics, but a number of popular methods determine if a set of modified genes has a higher than expected overlap with genes known to function as part of a pathway (functional enrichment testing). Recently, researchers have started to apply such analyses in a new way: to ask if the data they are collecting show similar disruptions to biological functions compared to reference data. Examples include studying whether similar pathways are perturbed in smokers vs. users of e-cigarettes, or whether a new mouse model of schizophrenia is justified, based on its similarity in cytokine expression to a previously published model. However, there is a dearth of robust statistical methods for testing hypotheses related to these questions and most researchers resort to *ad hoc* approaches. In this work, we propose a statistical approach to answering such questions. First, we propose a statistic for measuring the degree of equivalent change in individual genes across different treatments. Using this statistic, we propose applying gene set enrichment analysis to identify pathways enriched in genes that are affected in similar or opposing ways across treatments. We evaluate this approach in comparison to *ad hoc* methods on a simulated dataset, as well as two biological datasets and show that it provides robust results.

## Introduction

*In silico* functional genomics have become a standard approach in enabling researchers to use transcriptomics to understand biological pathways or molecular functions affected by the treatments or conditions they are studying. For example, a published protocol for the DAVID Bioinformatics Resources has over 15,000 citations(Huang et al. 2009). A paper discussing another popular approach, called Gene Set Enrichment Analysis (GSEA), has been cited over 13,000 times(Subramanian et al. 2005). Most methods determine if a set of modified genes has a higher than expected overlap with genes known to function as part of a pathway (functional enrichment testing)(Subramanian et al. 2005; Huang et al. 2009; Luo et al. 2009). Perhaps the simplest way of doing such an analysis is to perform overrepresentation analysis (ORA) and test if a set of genes (perhaps those that were statistically significantly differentially expressed between the comparator groups) overlaps a list of genes in a biological pathway more than what would be expected by chance(Huang et al. 2009),(Wang et al. 2017),(Kuleshov et al. 2016). Several important annotations of biological processes or pathways have been developed to facilitate such analyses. One of the best known is the Gene Ontology(Ashburner et al. 2000) (GO), although The Kyoto Encyclopedia of Genes and Genomes (KEGG)(Kanehisa et al. 2017) and Reactome(Fabregat et al. 2018) have additional annotations for the relationships between genes in a pathway. The statistical significance of overrepresentation of a significant set of genes in the genes of a pathway can be tested simply using the hypergeometric test. Other methods have been developed to improve the power and reliability of these tests, including GSEA(Subramanian et al. 2005). GSEA identifies sets of genes that group together near the top or bottom of a list of genes ranked by degree of differential expression (typically log_2_-fold change) more than one would expect by chance. There is no requirement that individual genes be statistically significant by whatever metric is used.

Now, some researchers are asking a different version of this question: i.e., they want to know if the data they are collecting show similar functional disruptions as compared to some reference data. For example, Shen, et al. studied whether similar pathways were perturbed in smokers vs. users of e-cigarettes(Shen et al. 2016). Gil-Pisa, et al. justified the use of their mouse model of schizophrenia based on its similarity in cytokine expression to a previously published model(Gil-Pisa et al. 2014). Martins-de-Souza, et al. showed that responders showed the same pathways were affected, but in opposite directions, in poor vs. good responders to anti-psychotics(Martins-de-Souza et al. 2015). Clearly, this idea has many potential applications. Unfortunately, we currently lack statistically sound approaches for most such analyses.

One possible approach is perform enrichment analysis separately for genes that are up and down regulated in each treatment and then find the intersection of pathways that move in the same or opposite directions(Ibanez et al. 2014; Sanchez-Valle et al. 2017). GSEA can test for pathways that are significantly up or down regulated. However, typically, researchers using this or similar approaches do not attempt to determine the probability of this occurring by chance, bringing the interpretability and reproducibility of such results into question. Also, these types of approaches do not indicate the degree to which pathways are changed in similar or opposing ways. We are not aware of any methods that can specifically address these questions. Furthermore, a substantial limitation of similar approaches is the underlying assumption that biological pathways depend on the co-expression of genes in them. Some pathways likely function in this manner, and one may be able to detect sub-pathways with equivalent or inverse changes, but the results will likely be biased to simple pathways.

For the case of drug-repurposing, Connectivity Mapping was introduced to address the need for detecting when genes are disrupted in similar ways. This approach assesses the correlation in ranked lists of genes, with the intent of identifying gene profiles for drugs that are correlated, or anti-correlated to a researcher’s own gene signature(Lamb et al. 2006), but this approach is not designed to identify specific biological functions that are similar across experiments. The same is true of other methods, such as openSesame(Gower et al. 2011) or the extreme cosine method (XCos)(Cheng et al. 2013), which were developed later. Even for drug repurposing, this may be an important point. That is, it may be important to be similar only in terms of certain pathways but not others. Therefore, a method that can systematically identify pathways with similar (or inverted) perturbations could be of great use. In this work, we propose a novel functional genomics approach called Equivalent Change Enrichment Analysis (ECEA) that seeks to accomplish this goal. We further introduce a novel metric called the Equivalent Change Index (ECI), which plays a key role in our proposed methodology. There are many potential applications of the proposed methodology, including the ability to focus on genes that may be more directly relevant to the experimental question, drug screening for treatments that have similar effects, demonstrating the viability of a new mouse model, and many other potential uses.

## Results

We present two key developments for identifying biological pathways that exhibit equivalent or inverse changes across experiments and/or treatments: i) the Equivalent Change Index (ECI), and ii) Equivalent Change Enrichment Analysis (ECEA). The ECI is a measure that is calculated at the level of individual genes from gene expression assays that ranges from [-1,1]. A value of 1 indicates that a gene was changed to the same degree by both treatments (e.g. a 2-fold change by each treatment compared to its respective control). A value of −1 indicates a gene was change in completely the opposite way (e.g. downregulated 2-fold by one treatment and upregulated 2-fold by the other). ECEA is a functional genomics approach that identifies pathways with a non-random distribution of equivalently or inversely changed genes.

We evaluated our approach on three datasets: one simulated, one biological data set with expected inverse changes, and a second biological data set with expected equivalent changes.

We benchmarked ECEA to the current approach that involves simply performing pathway enrichment analysis on the datasets from each treatment separately and then intersecting the results. We used both Gene Set Enrichment Analysis (GSEA) and over-representation analysis (ORA) for comparison. With GSEA, it is possible to get some idea of equivalent or inverse changes as well, because it tests for enrichment of up or downregulated genes.

### Simulations

The results of the simulations are shown for equivalent change in Fig. 1, and for inverse change in Fig. 2, using a probability of equivalent change or inverse change of 0.5 (i.e., each gene in a pathway chosen to have equivalent differential expression would have a probability of 0.5 of that equivalent change). Results at other probability levels are shown in the supplemental material. We did 100 simulations for each set of parameters. Each simulation involved 1 equivalently changed pathway, 1 inversely changed pathway, 1 pathway with differentially expressed genes but no relationship between experiments, and 7 pathways not affected by these simulated treatments. In each case, there were 5 samples for each treatment and 5 controls (N=20). The results show the proportion of times the equivalent or inversely changed pathway was detected. The inversely changed pathway was more difficult to detect by GSEA, because creating inverse changes will also tend to increase the number of genes that are not regulated in the same direction, which is a limitation of GSEA. For most levels of probability of differential expression and across levels of symmetry, ECEA outperformed GSEA or ORA. For equivalently changed pathways, GSEA outperformed ECEA only for low levels of probability of differential expression (PDE) and when the symmetry was extreme. In Fig. 3, the false positive rate (FPR) for detecting the pathway with differential expression but without enforced equivalent or inverse change is shown. For all levels of probability of differential expression ORA had the lowest FPR, but it is also very low for ECEA. When the symmetry is .1 or .9 the FPR is nearly linear for FGSEA, meaning that the greater the probability of differential expression, the greater the likelihood of identifying a pathway as having equivalent change by chance, using this approach.

**Figure 1:**
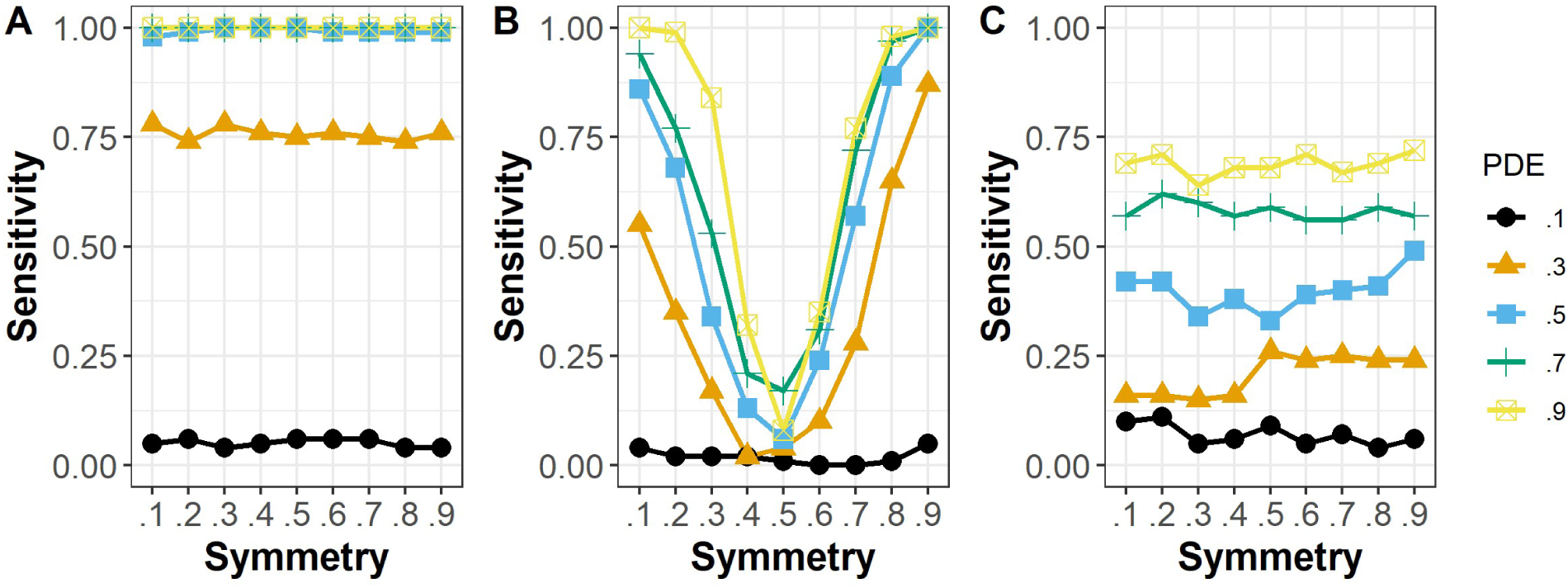
The proportion of equivalently changed pathways detected by each method, when the probability of equivalent change was 0.5. The x-axis displays the symmetry, which shows the probability of genes being up-regulated in the pathway as opposed to down-regulated (when they were differentially expressed). With a symmetry of 0.5, approximately half of the differentially expressed genes would be up-regulated and the rest would be down-regulated. The different lines show the sensitivity for different levels of probability of a gene being differentially expressed. The results are shown for (A) ECEA, (B) GSEA, and (C) ORA. For nearly all levels of probability of differential expression, ECEA was more sensitive than ORA. ECEA was also more sensitive than GSEA whenever the genes were not highly co-expressed (symmetry near .1 or .9).

**Figure 2:**
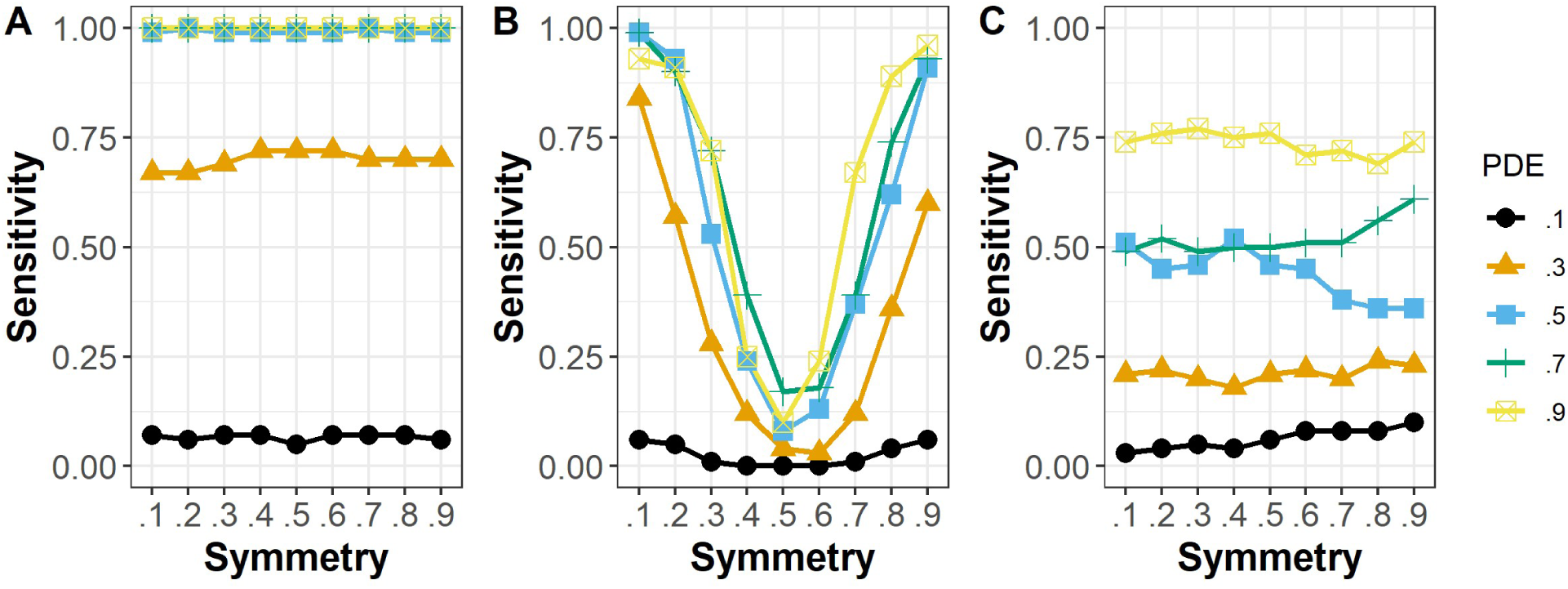
The proportion of inversely changed pathways detected by each method, when the probability of inverse change was 0.5. The x-axis displays the symmetry, which shows the probability of genes being up-regulated in the pathway as opposed to down-regulated (when they were differentially expressed). With a symmetry of 0.5, approximately half of the differentially expressed genes would be up-regulated and the rest would be down-regulated. The different lines show the sensitivity for different levels of probability of a gene being differentially expressed. The results are shown for (A) ECEA, (B) GSEA, and (C) ORA. For nearly all levels of probability of differential expression, ECEA was the most sensitive. ECEA was also more sensitive than GSEA whenever the genes were not highly co-expressed (symmetry near .1 or .9).

**Figure 3:**
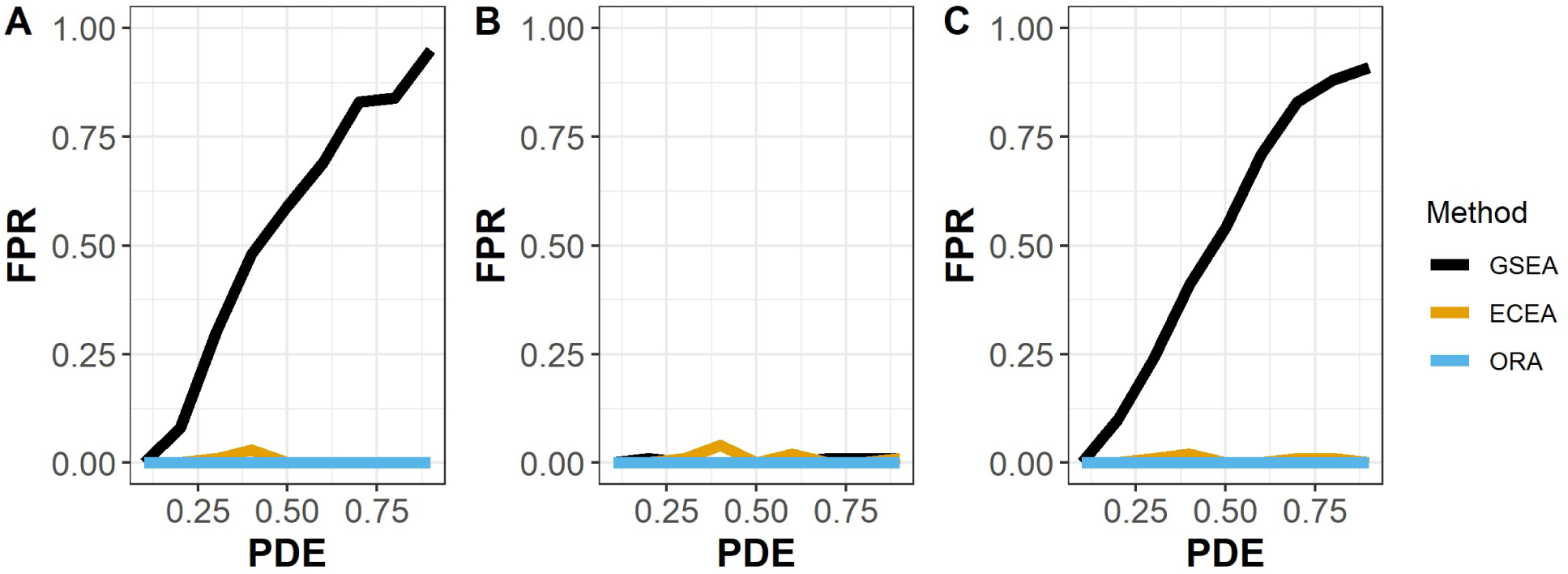
The proportion of times a pathway with differentially expressed genes was erroneously identified as being equivalently or inversely enriched by method with probability of equivalent change at 0.5. The sub-figures show the results at (A) Symmetry = 0.1, (B) Symmetry = 0.5, and (C) Symmetry = 0.9.

### *Glut4* Data

ECEA was run on the *Glut4* data to determine pathways enriched for genes that are equivalently or inversely changed when *Glut4* is knocked out or overexpressed in mice. The data were collected to determine the effect of *Glut4* on insulin sensitivity in white adipose tissue. The log_2_ fold change for each treatment vs. its respective control was calculated using the limma package for R(Ritchie et al. 2015). The log_2_ fold change was used to calculate the ECI, which was then used to perform the ECEA. First, we performed this analysis on the Kyoto Encyclopedia of Genes and Genomes (KEGG), with a false discovery rate cut-off of 0.25, which is the recommended threshold for GSEA(Subramanian et al. 2005). We found enrichment in equivalent or inverse change for 8 pathways using this approach. Of these, 5 were enriched with inversely changed genes and are listed in Table 1. It is important to note that the number of pathways equivalently or inversely changed do not represent a picture of overall equivalent or inverse change as pathways have many overlapping genes. For each enriched pathway, we have included the top 5 genes in the results, which are the genes with the greatest inverse change for the respective pathway.

**Table 1.** ECEA identified inversely changed KEGG pathways in the Glut4 data

| Pathway | FDR | NES | Size | Top 5 Genes |
| --- | --- | --- | --- | --- |
| mmu00280 Valine, leucine and isoleucine degradation | $1.85 \times 10^{-2}$ | -2.08 | 34 | Bckdha,Pccb,Acaa2,Mcee,Hmgcs1 |
| mmu00071 Fatty acid metabolism | $2.44 \times 10^{-1}$ | -1.62 | 32 | Acaa2,Eci1,Echs1,Acsl4,Acsl1 |
| mmu00640 Propanoate metabolism | $3.24 \times 10^{-2}$ | -1.83 | 21 | Pccb,Mcee,Echs1,Aldh7a1,Aldh1a1 |
| Mmu04080 Neuroactive ligand-receptor interaction | $2.44 \times 10^{-1}$ | -1.38 | 177 | Tshr,Sstr1,Pth1r,Vipr2,Fpr1 |
| mmu00310 Lysine degradation | $2.44 \times 10^{-1}$ | -1.64 | 25 | Echs1,Suv39h2,Aldh7a1,Aldh1a1,Ehmt2 |

Next, we applied the GSEA intersection approach. This involves finding pathway enriched for differentially expressed genes in each experiment separately and then intersecting the results. Using GSEA, for the *Glut4* KO, there were 32 significantly enriched pathways. For the overexpressed data, there were 29. One way to search for inversely changed pathways would be to find those enriched in upregulated genes using GSEA in one dataset and enriched for downregulated genes in the other (for equivalent changes we would simply intersect those changed in the same direction). Between these two results, there were 11 shared pathways. Of these, 6 had an inverse relationship and 5 and an equivalently changed relationship. Using this approach there is no way to assess the statistical significance of this relationship, but the pathways with inverse regulation are shown in Table 2.

**Table 2.** GSEA identified inversely changed KEGG pathways in the Glut4 data

| Pathway |
| --- |
| mmu00280 Valine, leucine and isoleucine degradation |
| mmu00071 Fatty acid metabolism |
| mmu03320 PPAR signaling pathway |
| mmu00640 Propanoate metabolism |
| mmu04146 Peroxisome |
| mmu04610 Complement and coagulation cascades |

Three of the pathways identified as being inversely regulated by ECEA were also identified by this GSEA approach. However, the total number of pathways available in the KEGG database is relatively limited at 225. Therefore, we also tried these approaches using the larger Reactome database, which has 1647 total pathways, which might allow for a more granular picture. Using ECEA, we found 27 pathways with significant enrichment, eight of which were inversely enriched (Table 3).

**Table 3.** ECEA identified inversely changed Reactome pathways in the Glut4 data

| Pathways | FDR | NES | Size | Top 5 Genes |
| --- | --- | --- | --- | --- |
| GPCR ligand binding | $1.72 \times 10^{-1}$ | -1.40 | 219 | Tshr,Ccr7,Ece1,Sstr1,Pth1r |
| Protein localization | $2.29 \times 10^{-1}$ | -1.52 | 68 | Dhrs4,Gstk1,Ech1,Slc25a17,Tysnd1 |
| SLC-mediated transmembrane transport | $6.69 \times 10^{-2}$ | -1.57 | 108 | Slc26a2,Slc31a1,Slc2a4,Slco3a1,Slc39a8 |
| Transport of inorganic cations/anions and amino acids/oligopeptides | $7.00 \times 10^{-2}$ | -1.69 | 43 | Slc26a2,Calm2,Slc7a8,Slc3a2,Slc4a8 |
| Peroxisomal protein import | $1.15 \times 10^{-1}$ | -1.68 | 40 | Dhrs4,Gstk1,Ech1,Tysnd1,Pex5 |
| VEGFR2 mediated vascular permeability | $1.29 \times 10^{-1}$ | -1.73 | 21 | Akt2,Calm2,Pak2,Ctnnb1,Nos3 |
| Transcriptional Regulation by E2F6 | $4.01 \times 10^{-2}$ | -1.79 | 15 | Phc1,Rbbp4,Ezh2,Ehmt2,Suz12 |
| SHC-mediated cascade:FGFR2 | $2.03 \times 10^{-1}$ | -1.68 | 17 | Fgf4,Fgf8,Grb2,Fgf6,Fgf18 |

Using the GSEA intersection approach, we found 14 pathways with inverse changes in regulation, two of which was also found by ECEA. These are shown in Table 4.

**Table 4.** GSEA identified inversely changed Reactome pathways in the Glut4 data

| Pathway |
| --- |
| Plasma lipoprotein assembly, remodeling, and clearance |
| Condensation of Prophase Chromosomes |
| Metabolism of vitamins and cofactors |
| Protein localization |
| Glucocorticoid biosynthesis |
| Hemostasis |
| Platelet activation, signaling and aggregation |
| Peroxisomal protein import |
| Platelet degranulation |
| Response to elevated platelet cytosolic Ca<sup>2+</sup> |
| Branched-chain amino acid catabolism |
| Regulation of Tp53 Degradation |
| Regulation of Tp53 Expression and Degradation |
| Laminin interactions |

An additional 31 pathways were identified using GSEA with the intersection approach that were enriched for differential expression with equivalent directional change in regulation.

It is worth noting that ECEA looks for enrichment in equivalent or inverse changes in the same genes across treatments, while the intersection approach will simply find overall changes in genes in the pathway that tend to be in the same direction. This might also be a useful thing to do, but the goals are slightly different. However, with the overlap approach, there is no indication as to whether the functional impact is likely to be similar, such as there is for ECEA (i.e. different parts of a complex pathway might be affected and thus not result in similar functional impacts).

Fig. 4 shows part of the VEGFR2 mediated vascular permeability pathway from Reactome. On the left, we can see the effect of the *Glut4* knockout compared to the controls, on the right the effect of *Glut4* overexpression. This pathway was identified by ECEA as inversely changed but not by GSEA. The figure illustrates a possible explanation why, due to the assumptions in GSEA about co-expression. There are clear inverse changes in Nos3, Akt2, Calm2, and Pdpk1 but some of these genes are upregulated and some downregulated in each treatment.

**Figure 4:**
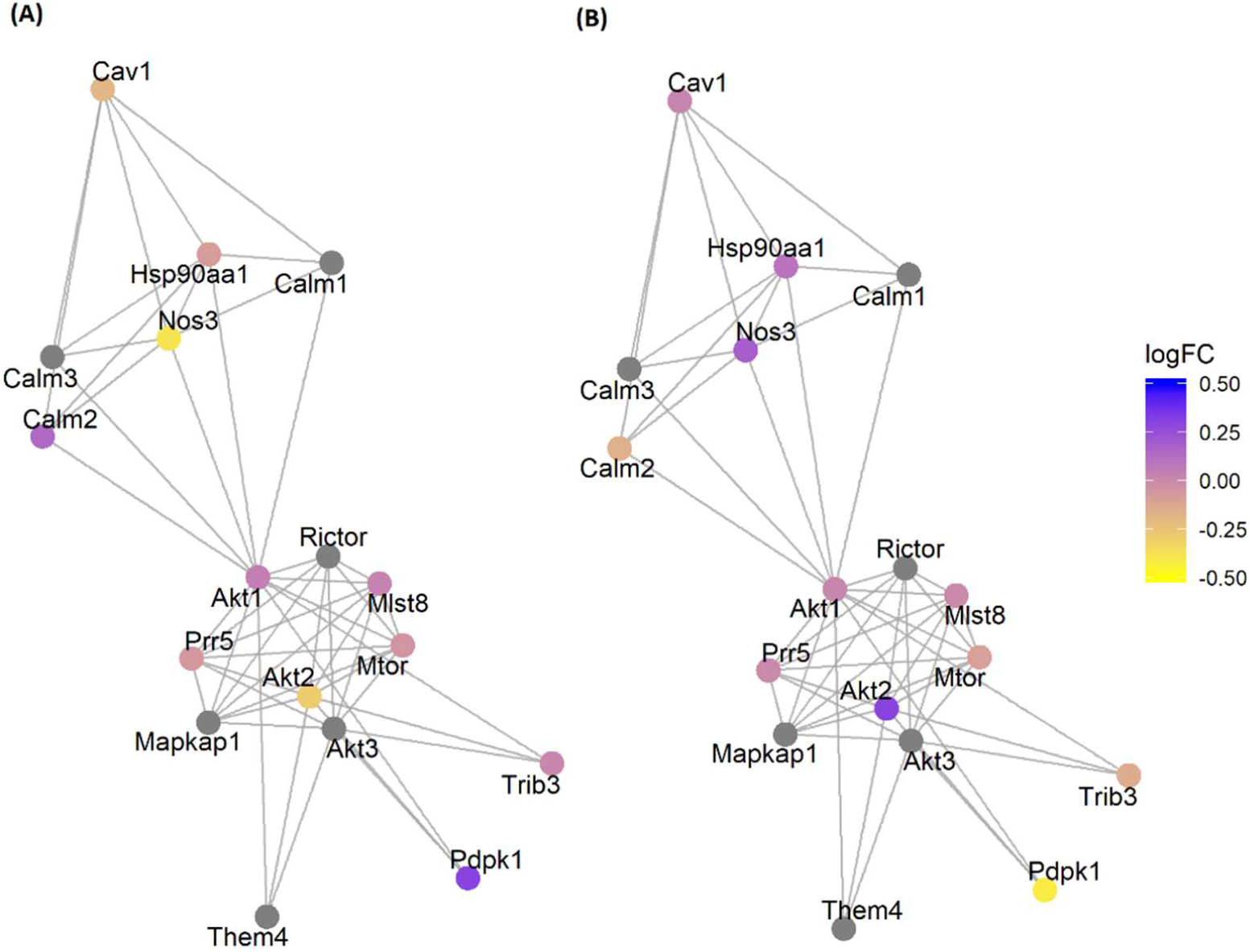
Differentially expressed genes in part of the VEGFR mediated vascular permeability pathway from Reactome. There are clear inverse changes between the two treatments pictured and, importantly, not all genes with inverse changes show the same direction in regulation (some are upregulated and others downregulated). (A) The log_2_-fold change in expression between Glut4 knockout and control. (B) The log_2_-fold change in expression between Glut4 overexpressed and control.

### Antidepressant Data

We analyzed the antidepressant data in much the same way as the *Glut4* data, except in these data we expected to capture equivalent changes. These data were collected to investigate the effect of two different antidepressant drugs, ketamine and imipramine, on a mouse model of depression. For the KEGG pathways, in this case, there were 6 pathways with significant enrichment for equivalent change across treatments (Table 5) and none for inverse change.

**Table 5.** ECEA identified equivalently changed KEGG pathways in the antidepressant data

| Pathway | FDR | NES | Size | Top 5 Genes |
| --- | --- | --- | --- | --- |
| mmu03010 Ribosome | 1.01x10<sup>-2</sup> | 1.52 | 95 | Rpl7a,Rpsa,Rpl34,mt-Rnr1,Rplp1 |
| mmu00230 Purine metabolism | 2.12x10<sup>-1</sup> | 1.22 | 144 | Ada,Nt5e,Adcy4,Ak7,Pde1c |
| mmu00500 Starch and sucrose metabolism | 2.12x10<sup>-1</sup> | 1.47 | 22 | Gaa,Amy1,Gys1,Pgm1,Pgm2 |
| mmu00250 Alanine, aspartate and glutamate metabolism | 2.12x10<sup>-1</sup> | 1.44 | 28 | Gad2,Nit2,Aldh4a1,Gad1,Ass1 |
| mmu04080 Neuroactive ligand-receptor interaction | 5.36x10<sup>-2</sup> | 1.24 | 185 | Mc3r,Sstr5,Crhr2,Glp1r,Gabrq |
| mmu00340 Histidine metabolism | 1.01x10<sup>-2</sup> | 1.67 | 20 | Aldh3b1,Aldh7a1,Hdc,Aldh3a1,Ddc |

When we performed the GSEA intersection approach, we found 2 pathways that were significantly enriched after treatment with either drug and both had changes in the same direction (Table 6). Both pathways were also identified by the ECEA approach.

**Table 6.**
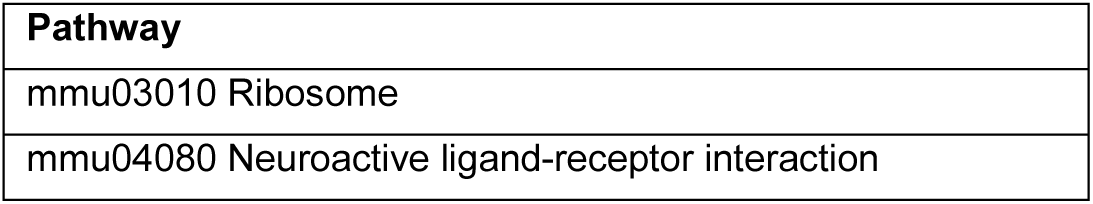
GSEA identified equivalently changed KEGG pathways in the antidepressant data

Next, we applied ECEA, using these data, to the Reactome database. The results are shown in Table 7. A total of 17 pathways were found to be enriched for equivalent changes across the two experiments and none for inverse changes.

**Table 7.** ECEA identified equivalently changed Reactome pathways in the antidepressant data

| Pathway | FDR | NES | Size | Top 5 Genes |
| --- | --- | --- | --- | --- |
| Regulation of Insulin-like Growth Factor (IGF) transport and uptake by Insulin-like Growth Factor Binding Proteins (IGFBPs) | $2.76 \times 10^{-2}$ | 1.38 | 78 | Gpc3,Trf,Scg3,Penk,F5 |
| Post-translational protein phosphorylation | $3.02 \times 10^{-2}$ | 1.38 | 73 | Gpc3,Trf,Scg3,Penk,F5 |
| GPCR ligand binding | $1.14 \times 10^{-1}$ | 1.22 | 200 | Mc3r,Sstr5,Crhr2,Glp1r,Nts |
| L13a-mediated translational silencing of Ceruloplasmin expression | $8.07 \times 10^{-3}$ | 1.39 | 100 | Rpsa,Rpl34,Rplp1,Rps29,Rpl8 |
| Eukaryotic Translation Initiation | $8.07 \times 10^{-3}$ | 1.35 | 108 | Rpsa,Rpl34,Rplp1,Rps29,Rpl8 |
| Formation of a pool of free 40S subunits | $8.07 \times 10^{-3}$ | 1.45 | 90 | Rpsa,Rpl34,Rplp1,Rps29,Rpl8 |
| GTP hydrolysis and joining of the 60S ribosomal subunit | $1.48 \times 10^{-2}$ | 1.38 | 101 | Rpsa,Rpl34,Rplp1,Rps29,Rpl8 |
| Cap-dependent Translation Initiation | $8.07 \times 10^{-3}$ | 1.35 | 108 | Rpsa,Rpl34,Rplp1,Rps29,Rpl8 |
| Translation | $3.02 \times 10^{-2}$ | 1.23 | 211 | Mrpl10,Mrps16,Rpsa,Rpl34,Rplp1 |
| SRP-dependent cotranslational protein targeting to membrane | $8.07 \times 10^{-3}$ | 1.53 | 81 | Rpsa,Rpl34,Rplp1,Rps29,Rpl8 |
| Major pathway of rRNA processing in the nucleolus and cytosol | $8.07 \times 10^{-3}$ | 1.36 | 156 | Exosc10,Rpsa,Rpl34,Rplp1,Rps29 |
| rRNA processing | $8.07 \times 10^{-3}$ | 1.36 | 156 | Exosc10,Rpsa,Rpl34,Rplp1,Rps29 |
| rRNA processing in the nucleus and cytosol | $8.07 \times 10^{-3}$ | 1.36 | 156 | Exosc10,Rpsa,Rpl34,Rplp1,Rps29 |
| Nonsense-Mediated Decay (NMD) | $8.07 \times 10^{-3}$ | 1.44 | 102 | Upf2,Rpsa,Rpl34,Rplp1,Rps29 |
| Nonsense Mediated Decay (NMD) independent of the Exon Junction Complex (EJC) | $8.07 \times 10^{-3}$ | 1.51 | 83 | Rpsa,Rpl34,Rplp1,Rps29,Rpl8 |
| Nonsense Mediated Decay (NMD) enhanced by the Exon Junction Complex (EJC) | $8.07 \times 10^{-3}$ | 1.44 | 102 | Upf2,Rpsa,Rpl34,Rplp1,Rps29 |
| Sulfur amino acid metabolism | $8.07 \times 10^{-3}$ | 1.74 | 19 | Slc25a10,Gm4737,Ahcy,Cdo1,Adi1 |

Finally, we used the GSEA intersection approach to examine these same data. A total of 20 pathways were found to be enriched for differentially expressed genes and regulated in the same direction by both drugs. These are shown in Table 8. Eight of these pathways were also identified by ECEA.

**Table 8.** GSEA identified equivalently changed pathways in the antidepressant data

| Pathway |
| --- |
| Signaling by GPCR |
| Class A/1 (Rhodopsin-like receptors) |
| Peptide ligand-binding receptors |
| GPCR downstream signaling |
| G alpha (i) signalling events |
| GPCR ligand binding |
| L13a-mediated translational silencing of Ceruloplasmin expression |
| Formation of a pool of free 40S subunits |
| GTP hydrolysis and joining of the 60S ribosomal subunit |
| G alpha (q) signalling events |
| G alpha (s) signalling events |
| SRP-dependent cotranslational protein targeting to membrane |
| Nonsense-Mediated Decay (NMD) |
| Nonsense Mediated Decay (NMD) independent of the Exon Junction Complex (EJC) |
| Nonsense Mediated Decay (NMD) enhanced by the Exon Junction Complex (EJC) |
| Protein-protein interactions at synapses |
| Dopamine Neurotransmitter Release Cycle |
| Serotonin Neurotransmitter Release Cycle |
| Glutamate Neurotransmitter Release Cycle |
| Synaptic adhesion-like molecules |

## Discussion

In this work we have presented a new approach to functional genomic analysis that can identify key biological pathways that are changed in similar (equivalent) or opposing (inverse) ways across diverse experiments. Due to our unique approach, data collected at different times, by different groups can be used, because there is no comparison of gene expression values, just the effect sizes, and there is no dependence on exact estimates of those effects. This has the potential to allow researchers to capitalize on publicly available data in new ways. However, it works just as well when two treatments are run as part of the same project. Equivalent change enrichment analysis (ECEA) allows researchers to determine the specific pathways and functions that are regulated in close to the same fashion by different treatments, or pathways for which changes can be reversed by one treatment compared to another.

As we demonstrate, a similar type of analysis can be done by intersecting the results of two separate enrichment analyses, and this is undoubtedly a useful technique. However, such approaches cannot determine if a pathway is changed in the same way (i.e. the same parts of the pathway) or to the same extent. For larger pathways, the differences may be critical.

On the simulation data, all approaches were able to capture a useful proportion of equivalently and inversely changed pathways in most situations. However, ECEA typically had the most consistent performance. Also, for the GSEA intersection approach, a number of its results are likely to be false positives, particularly if there are many genes that are differentially expressed. For pathways with mostly co-expressed genes, the GSEA intersection approach may have somewhat more power than ECEA, however, this approach will be unable to identify pathways with equivalent or inverse changes when there are genes that are both up and down-regulated by a treatment. It is important to note that this is not a limitation of GSEA, given it is its intended behavior. It is simply a limitation of applying GSEA in this context.

Our approach is invariant to the direction of change in gene expression, because we are determining enrichment in similar or inverse changes across experiments. Therefore, a change can be equivalent for multiple genes, even if they are up or down-regulated in the same pathway. Furthermore, our ECEA approach outlined here can calculate the statistical significance of enrichment in genes that are dysregulated in similar or opposing ways across experiments. This should be particularly useful when applied to larger pathways, because enrichment will only be found when the same parts of the pathway are affected similarly (rather than a similar trend in expression on average for the pathway overall).

One potential limitation of the ECEA approach is the somewhat restrictive assumption that equivalent change in a pathway means that the same genes change to the same degree by different treatments. This will undoubtedly be more or less useful of an assumption in different contexts and should be kept in mind when using our approach.

In the *Glut4* dataset, both ECEA and GSEA identified the “mmu00280 Valine, leucine and isoleucine degradation” KEGG pathway as one with inverse changes. Indeed, prior research has shown that increases in circulating branched chain amino acids are associated with insulin resistance in obese patients(Zhao et al. 2016), which is relevant for this dataset investigating the effect of *Glut4* on insulin sensitivity in white adipose tissue. ECEA also identified the “VEGFR2 mediated vascular permeability” Reactome pathway as having inverse changes across the treatments. Interestingly, this pathway is specifically related to the experimental question, as research has shown that VEGFR2 can modulate insulin sensitivity in white adipose tissue(Honek et al. 2014). However, the GSEA intersection approach identified the “Response to elevated platelet cytosolic Ca2+” Reactome pathway as having inverse changes, and Ca2+ has also been linked to regulation of insulin signaling(Kang et al. 2017). Thus, the ECEA approach has some statistical advantages, but there are specific circumstances for which the GSEA intersection approach will work well. It is seldom the case in functional genomics that a single method can be claimed to have every advantage. Also, our ECEA approach will identify when pathways are change in the same or specifically opposing ways (i.e. same genes) while the intersection approach will more generally identify pathways that experience overall changes in gene expression that are similar or opposing. Depending on one’s needs, this is a key difference that should be kept in mind. Nevertheless, these results indicate our approach can at least identify inversely changed pathways across treatments that are relevant to the target disease, and importantly, assign a statistical significance to the results.

In the antidepressant data, both ECEA and GSEA identified equivalent regulation by antidepressants of ribosome-related genes and indeed this association has been observed in patients with depression compared to healthy controls(Hori et al. 2018). This suggests that both ketamine and imipramine have similar influence on the regulation of genes involved that might serve as biomarkers of depression and suggest the utility of our approach in a precision medicine context. ECEA identified the “Regulation of Insulin-like Growth Factor (IGF) transport and uptake by Insulin-like Growth Factor Binding Proteins (IGFBPs)” Reactome pathway as equivalently changed by ketamine and imipramine and there is research linking this pathway with depression(Wang et al. 2014). However, the GSEA approach identified the “Glutamate Neurotransmitter Release Cycle” Reactome pathway as equivalently changed and this pathway has also been linked to depression(Duman et al. 2019). Thus, we can see that both approaches can lead to biologically meaningful, yet different, results. Although we highlight these specific results as examples, there are other pathways that seem to have a direct connection to both datasets identified by both methods.

The results for the antidepressant data are particularly exciting, because they demonstrate an important use case for our approach. Data for a new drug can be collected and commonalities in functional effects in comparison with existing drugs can be predicted, using a model organism. This has clear implications for the field of drug repositioning. One could imagine inverse enrichment could play a similarly important role, by allowing the prediction of a drug reversing the changes in genes of pathways disrupted in a disease. Although the GSEA intersection approach will work, the ECEA approach will particularly identify pathways where the changes are equivalent or inverted at the gene level within the pathway, which may be more useful when considering targeted treatments.

ECEA is not a general purpose functional genomics approach that will supplant existing computational functional genomics methods. Rather, we have demonstrated that it is a useful new tool that can allow researchers to garner relevant new insights into certain kinds of data. It allows for statistical rigor to be brought to research questions that are already being investigated in other ways, and potentially opens new avenues of inquiry.

## Methods

### Equivalent Change Index

Our goal is to be able to identify genomic pathways that are changed equivalently or inversely given two sets of experiments, each with a treatment and control. As a first step, we need a metric for the degree of equivalent change for a single gene. With such a metric, we can then search for pathways that have an unusual degree of equivalently or inversely changed genes. Therefore, we introduce the idea of an Equivalent Change Index (ECI).

Let 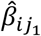 and 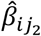 be the estimated effect sizes (ES) of two experiments for gene *i* on experiment *j*. The ES could be a log_2_ fold change, a standardized mean difference, or a simple mean difference. Then we define the equivalent change index (ECI) for gene *i* to be:

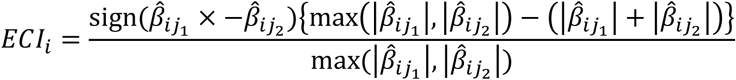

This is simply the ratio of the smaller ES to the larger ES in terms of absolute value of both effects and with a sign reflecting whether the effects were in the same direction (positive) or opposite directions (negative). Thus, *ECI*_*i*_ ∈ [−1,1]. An *ECI*_*i*_ of 1 means that the ES was exactly the same for gene *i* in both experiments. Likewise, an *ECI*_*i*_ of −1 means the ES was exactly opposite in both experiments. Therefore, *ECI*_*i*_ indicates either the degree of equivalence or inverseness for a gene in one experiment compared to a separate experiment, depending on its sign.

### Pathway-level Equivalent and Inverse Change

The ECI gives us a way to find pathways enriched for genes changed in equivalent or inverse ways across experiments. A high ECI indicates equivalent change, not the directionality of the change.

Therefore, if some genes in a pathway are up-regulated and others down-regulated by one treatment, but the reverse happens in a second treatment, the ECI will be low for all genes in the pathway. This is a critical property for our approach.

As an example, we can consider an experiment to determine the genomic influence of Drug B. In particular, we want to know biological functions for which Drug B has similar effects as Drug A. We measure gene expression for patients on Drug B and a control. We already have similar data for Drug A. Therefore, we calculate the ECI for Drug B and A on each gene. Next, we rank all genes by their ECI. We can now consider a hypothetical functional pathway, with a set of genes *P*. The full list of genes in the experiment is set *G*. The function *g*(*x*) yields the index of a gene in *G* with rank *x*, with genes ranked by ECI. We further define *S* = {*i*|*G*_*i*_ ∈ *P*} and *R* = {*i*|*G*_*i*_ ∉ *P*}, where *S* is the indices of the genes in the pathway, and *R* is the indices of the genes not in the pathway. Now, we will consider a gene of rank *x*. Suppose:

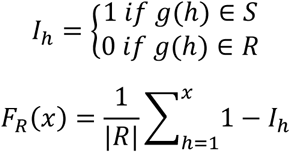

where |*R*| represents the number of genes not in the pathway under consideration. This gives us the proportion of genes that are not in the pathway and have a rank of *x* or less. We also have:

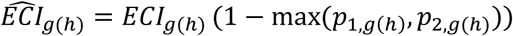

where *p*_1,*g*(*h*)_ and *p*_2,*g*(*h*)_ are the p-values for the test used to determine the effect sizes for *g*(*h*) in experiments 1 and 2. This yields a weighted ECI, based on the maximum p-value of the ES from each dataset for a given gene. This weight is useful to separate genes with a high or low ECI based on our confidence in their differential expression results.

Next, suppose:

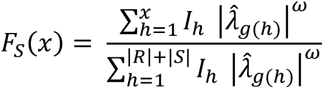

When *ω* = 0, this gives the proportion of genes that are in the pathway and have a rank of *x* or less. When *ω* ≠ 0, we will get a weighted ratio, depending on 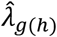, which we will discuss more in a minute.

Now, we can determine a quantity *D*:

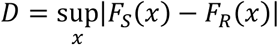

When *ω* = 0, this *D* is the Kolmogorov-Smirnov statistic. Therefore, we could use it for a hypothesis test of whether the distribution of ranks is different for a particular pathway vs. all other pathways. Unfortunately, this is problematic. The K-S test depends on an assumption of independence, which gene expression data cannot claim. The K-S test has been shown to be sensitive to violations of this assumption, so it is important to consider. Therefore, we will set *ω* = 1, which will provide a weighted Kolmogorov-Smirnov statistic. Note that this is exactly the approach taken by GSEA. The difference is, we have substituted the ECI for the local statistic used by GSEA (which is based on effect size)(Subramanian et al. 2005). Thus, *F*_*S*_(*x*) is adjusted for correlation between genes that is not associated with the treatments (this assumes that genes in a pathway would tend be correlated), because it is higher when the genes tend to be more equivalently expressed as the result of two treatments and is not a simple proportion. *D* is nevertheless still dependent on the size of the gene set. Therefore, we will scale *D*, getting 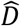, and perform permutation testing for enrichment in the same manner as GSEA, by using the fgsea package for R. We call this approach equivalent change enrichment analysis or ECEA.

One convenient aspect of the statistic 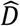, is that it represents a directional effect size for the entire pathway. Therefore, it can be used to judge the overall equivalent or inverse change of genes in a pathway, which is a particularly useful result for functional genomics. Unlike in standard GSEA, ECEA makes no assumption about the directionality of the change for gene effects. That is, genes that are up-regulated across treatments receive a high ECI, as do genes that are down-regulated across treatments. Therefore, there is no implicit assumption of co-expression for a pathway, meaning that ECEA can be used to investigate a wider array of pathways than typical GSEA, although for an entirely different use case.

### Data

In order to assess our method in both controlled and realistic conditions, we examined the performance of our approach using both simulated and biological data.

### Simulation

The simulated gene expression data were created using an approach similar to the one created by Dembele (2013)(Dembele 2013). This approach simulates correlation structure between genes, like might occur in a biological pathway in real data, and allows us to simulate different treatments that affect those simulated pathways to varying degrees. We modified the approach to create datasets in which genes perturbed by one treatment have a chance of being similarly perturbed (or inversely) by a second treatment.

We simulated 72,900 gene expression data sets, with correlation structure (the reason for this number is explained shortly). Each dataset was constructed from different runs of the algorithm, using different parameters for the proportion of equivalent or inversely changed genes and then combined, each subset thus representing a pathway, with its own correlation structure. Thus, we are making the simplifying assumption that there is no correlation between pathways in the simulation and there are no overlapping genes between pathways. For each dataset, we created seven pathways with no treatment effect (to provide a background), one pathway with equivalent change, one with inverse change, and finally a pathway without equivalent or inverse change but that was still enriched for differentially expressed genes. Varying the probability of genes being differentially expressed we ran 100 simulations at each probability level (from 0.1 to 0.9 in increments of 0.1). We also varied the symmetry of changes in a pathway (from 0.1 to 0.9 in increments of 0.1), and the probability of equivalent or inverse change (from 0.1 to 0.9). By symmetry, we mean proportion that were up vs down regulated. Thus, a symmetry of .5 means there was an equal probability of up vs. down regulation for differentially expressed genes. For each resulting dataset, we then calculated the number of times each equivalently or inversely changed pathway was detected by our method and calculated the number of true positives (TP), false positives (FP), true negatives (TN), and false negatives (FN), which were used to calculate other statistics, such as the sensitivity. We applied ECEA, GSEA, and ORA to these data and determined sensitivity and false positive rate for the recovery of pathways with varying levels of equivalent or inversely changed genes.

### Biological Data

The first biological dataset was created by Kraus, et al.(Kraus et al. 2014), to study the effect of the *Glut4* gene in adipose tissue on insulin sensitivity in a mouse model. These data were specifically created to have opposing effects and represent a dataset with expected inverse changes between two treatments. These samples are available in the National Center for Biotechnology Information’s (NCBI’s) Gene Expression Ominbus (GEO)(Edgar et al. 2002; Barrett et al. 2013) under accession GSE35378. This study involved 12 mice: 3 were adipose-*Glut4*-/-, 3 were aP2-Cre transgenic mice (controls for the *Glut4*-/-), 3 were adipose-*Glut4*-Tg mice with *Glut4* transgenically overexpressed, and 3 were FVB mice (controls for the adipose-*Glut4*-Tg mice). Gene expression was assayed using the Affymetrix Murine Genome U74A Version 2 Array. These were background subtracted and normalized using the rma function of the oligo package(Carvalho and Irizarry 2010) for the R statistical environment(Team 2017). Differential expression was assessed using the limma package(Ritchie et al. 2015) for R.

The second biological dataset was created by Bagot, et al.(Bagot et al. 2017) to investigate the effect of two antidepressants on the transcriptome in a mouse model of depression. Therefore, we expect to find some equivalent changes in this dataset. Gene expression was assayed using RNA sequencing on the Illumina HiSeq 2500 platform. Two drugs were examined, ketamine and imipramine, and various brain regions were examined. We limited our analysis to mice that were susceptible to depression and only used samples from the prefrontal cortex (PFC), in order to control confounding and because PFC had the greatest number of these samples available. Differential expression was assessed using the DESeq2(Love et al. 2014) package for R.

For the biological data, we do not have a ground truth. Here, our focus will be on determining whether our approach can detect equivalent or inverse changes that appear to make sense given the models and treatments. For each dataset, we will be examining the ability of ECEA and GSEA to identify disease relevant pathways.

## Availability

ECEA is available as a package for the R statistical environment, at https://github.com/jeffreyat/ECEA.

## Acknowledgements

This study was supported by the NIH 5P20GM130423 through the Kansas Institute of Precision Medicine and used the Quantitative ‘Omics Core (QOC).

