## supplemental material for "Equivalent Change Enrichment Analysis: Assessing Equivalent and Inverse Change in Biological Pathways between Diverse Experiments"

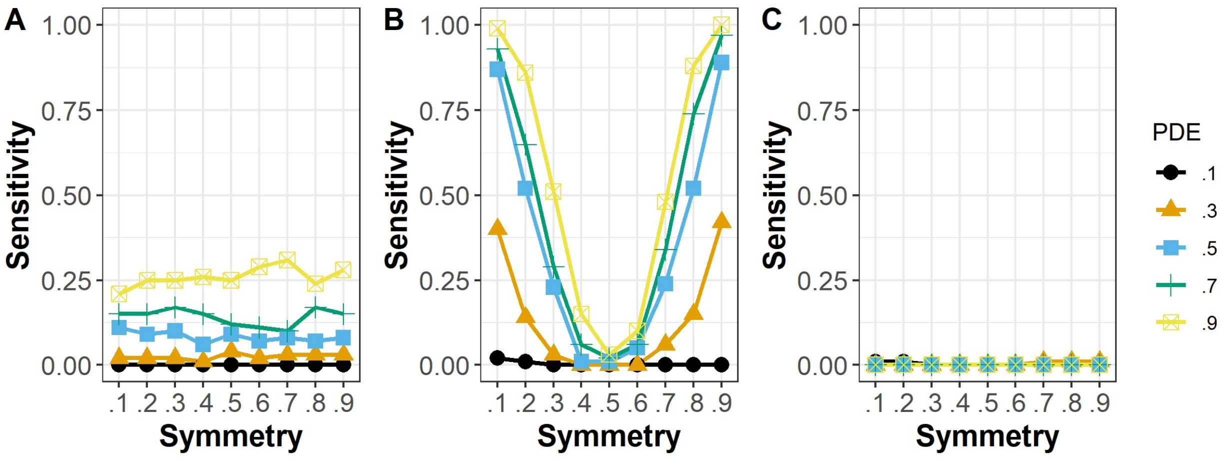


*Figure S1: The proportion of equivalently changed pathways detected by each method, when the probability of equivalent change was 0.1. The results are shown for (A) ECEA, (B) GSEA, and (C) ORA.*

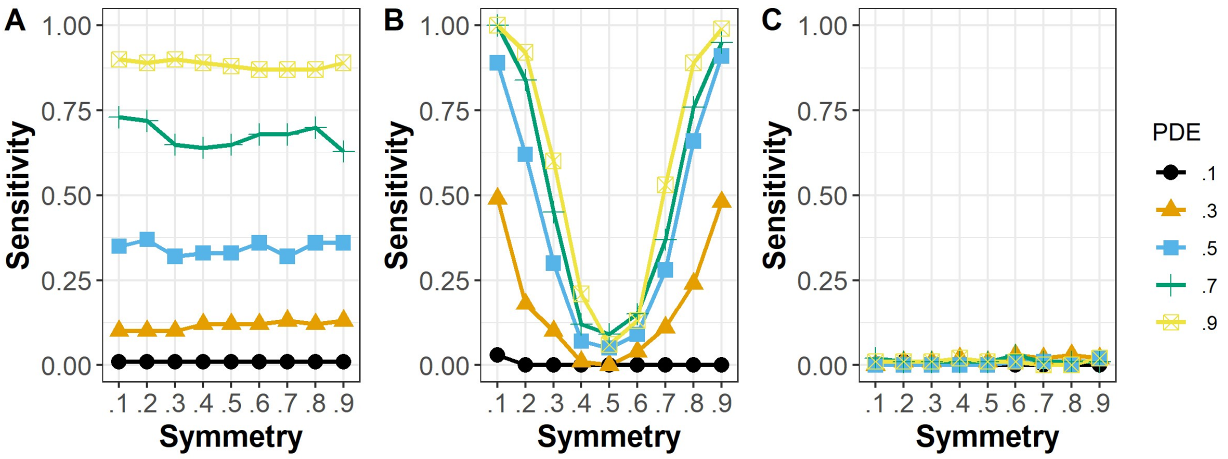


*Figure S2: The proportion of equivalently changed pathways detected by each method, when the probability of equivalent change was 0.2. The results are shown for (A) ECEA, (B) GSEA, and (C) ORA.*

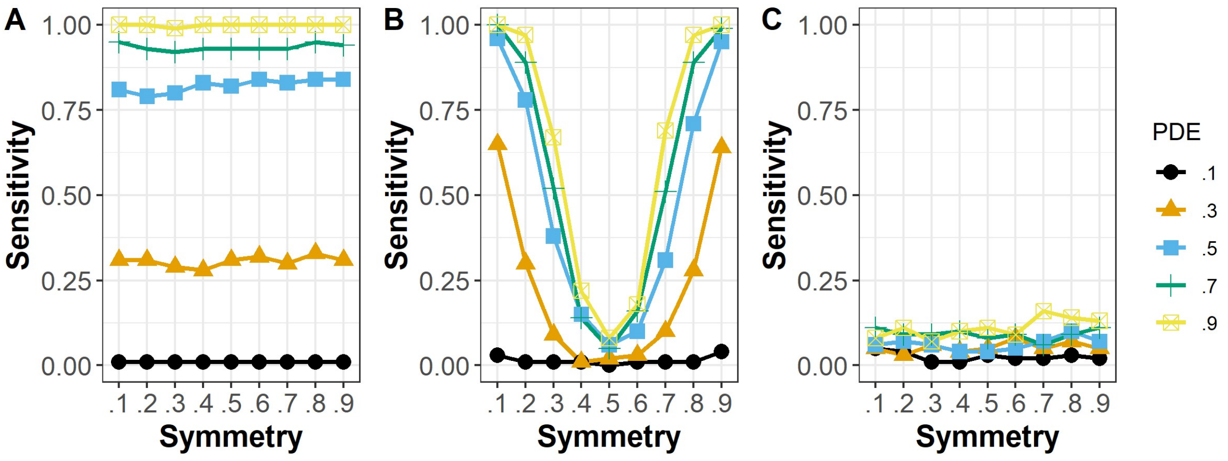

*Figure S3: The proportion of equivalently changed pathways detected by each method, when the probability of equivalent change was 0.3. The results are shown for (A) ECEA, (B) GSEA, and (C) ORA.*

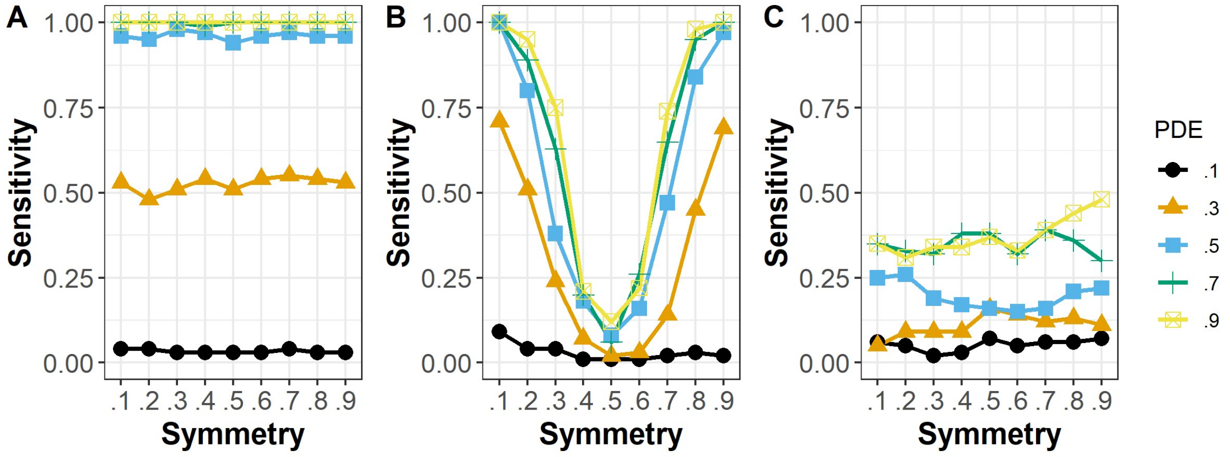


*Figure S4: The proportion of equivalently changed pathways detected by each method, when the probability of equivalent change was 0.4. The results are shown for (A) ECEA, (B) GSEA, and (C) ORA.*

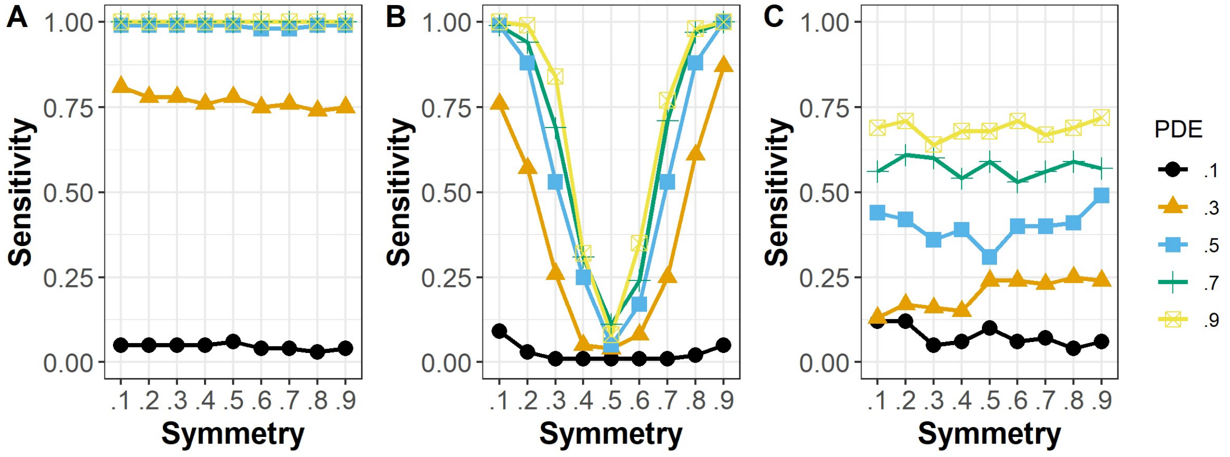


*Figure S5: The proportion of equivalently changed pathways detected by each method, when the probability of equivalent change was 0.5. The results are shown for (A) ECEA, (B) GSEA, and (C) ORA.*

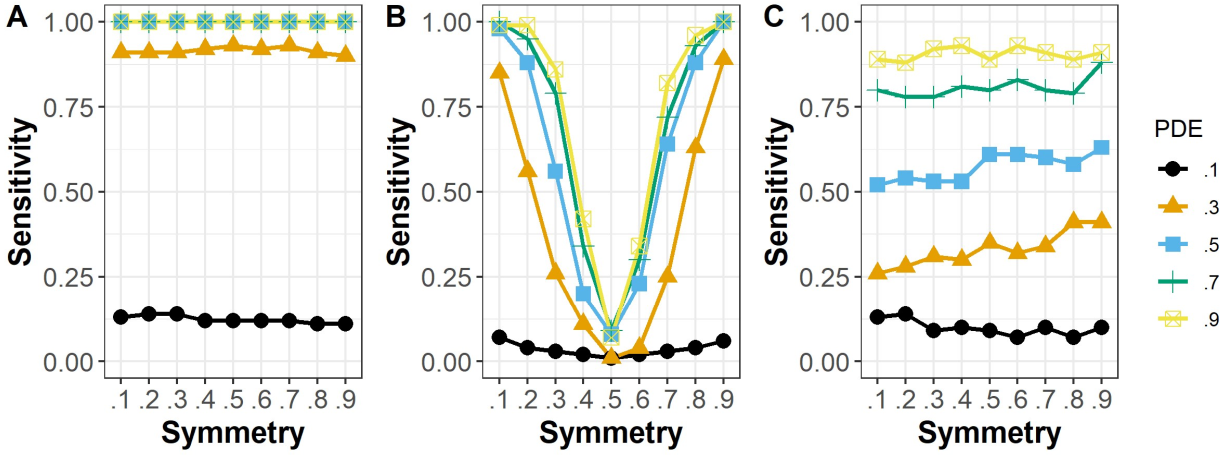

*Figure S6: The proportion of equivalently changed pathways detected by each method, when the probability of equivalent change was 0.6. The results are shown for (A) ECEA, (B) GSEA, and (C) ORA.*

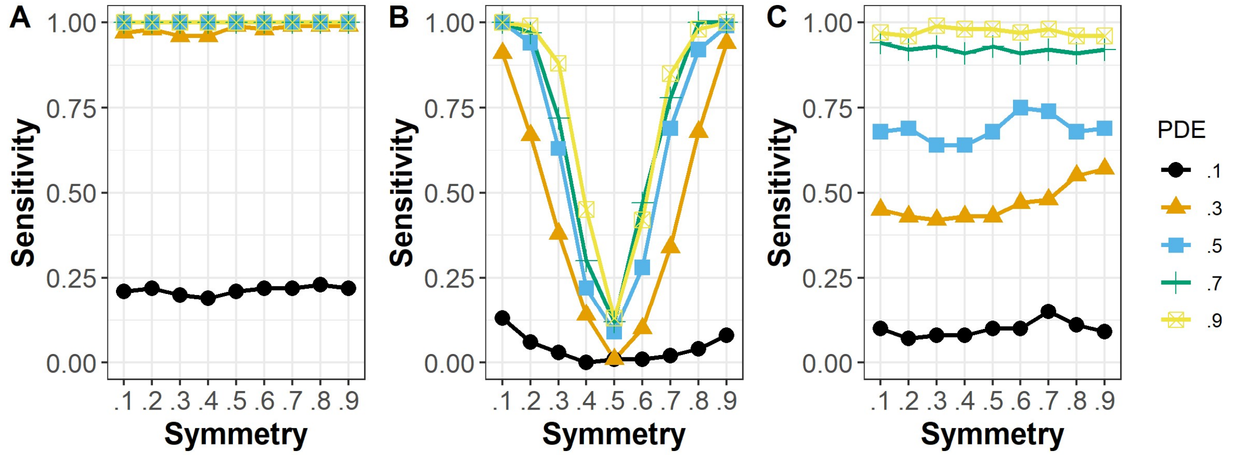


*Figure S7: The proportion of equivalently changed pathways detected by each method, when the probability of equivalent change was 0.7. The results are shown for (A) ECEA, (B) GSEA, and (C) ORA.*

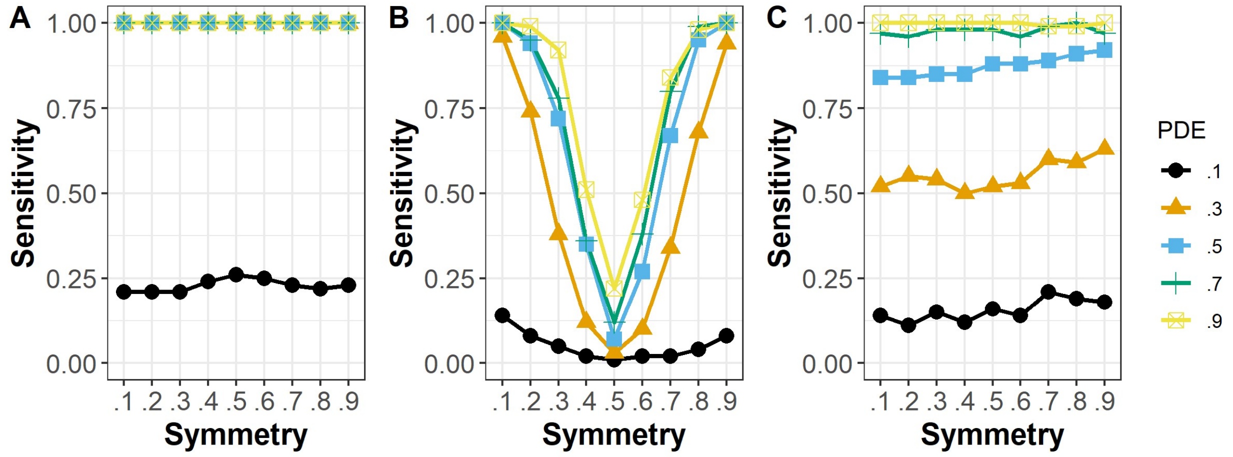


*Figure S8: The proportion of equivalently changed pathways detected by each method, when the probability of equivalent change was 0.8. The results are shown for (A) ECEA, (B) GSEA, and (C) ORA.*

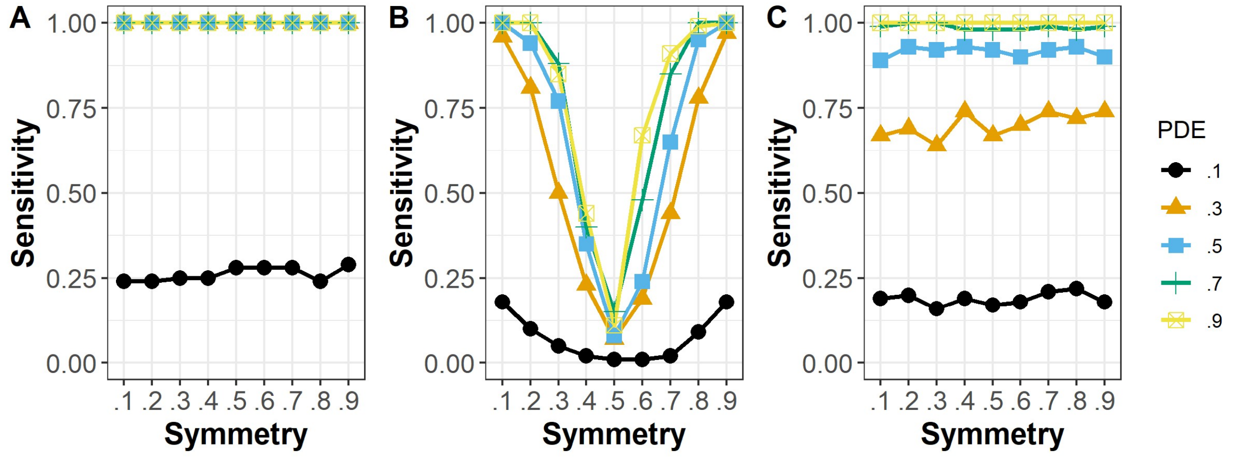

*Figure S9: The proportion of equivalently changed pathways detected by each method, when the probability of equivalent change was 0.9. The results are shown for (A) ECEA, (B) GSEA, and (C) ORA.*

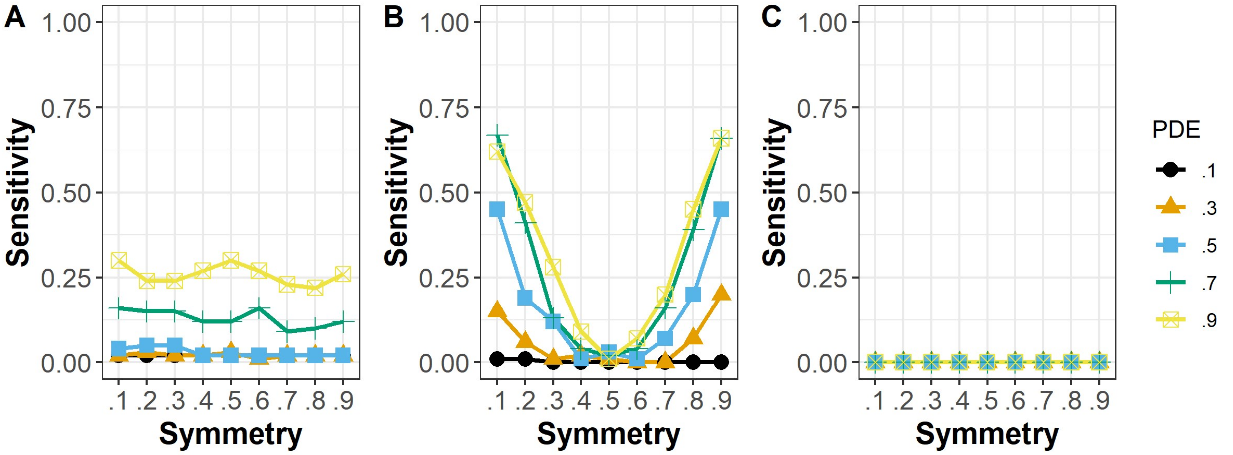


*Figure S20: The proportion of inversely changed pathways detected by each method, when the probability of inverse change was 0.1. The results are shown for (A) ECEA, (B) GSEA, and (C) ORA.*

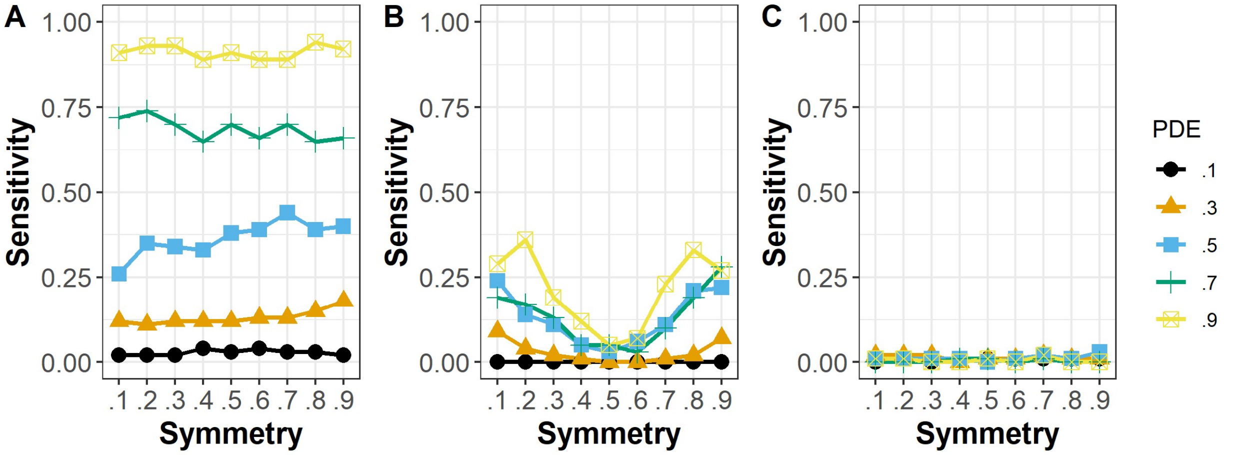


*Figure S31: The proportion of inversely changed pathways detected by each method, when the probability of inverse change was 0.2. The results are shown for (A) ECEA, (B) GSEA, and (C) ORA.*

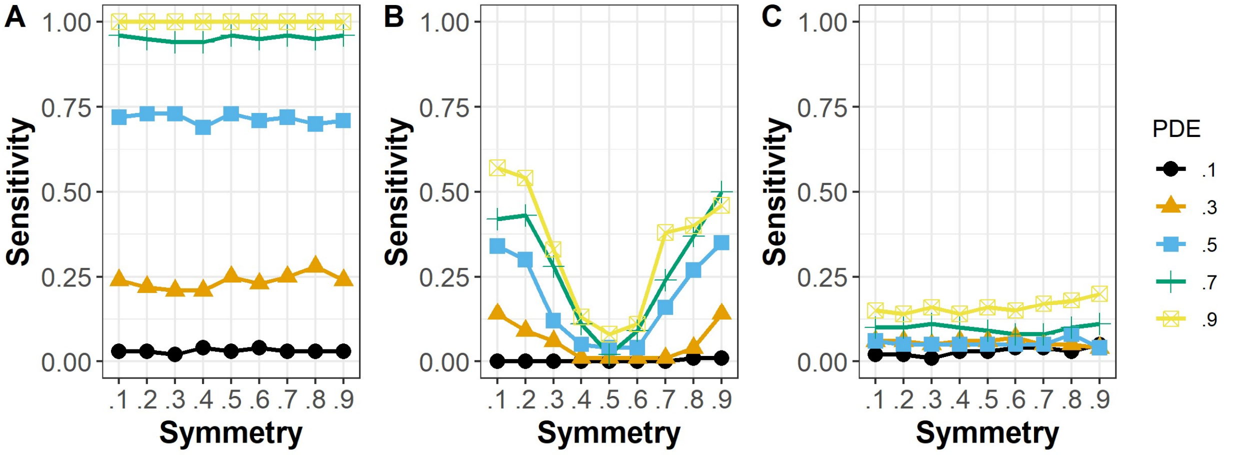

*Figure S42: The proportion of inversely changed pathways detected by each method, when the probability of inverse change was 0.3. The results are shown for (A) ECEA, (B) GSEA, and (C) ORA.*

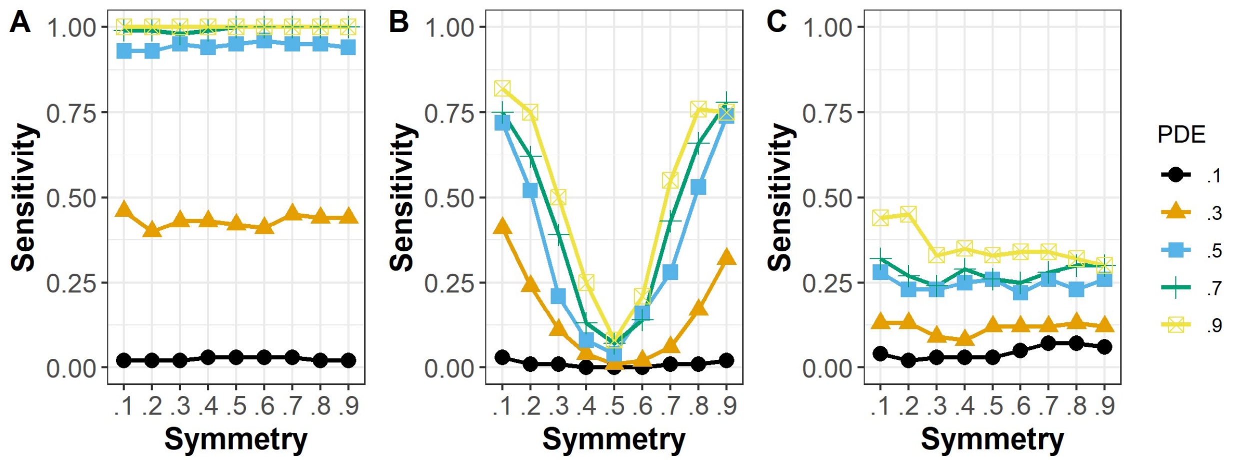


*Figure S53: The proportion of inversely changed pathways detected by each method, when the probability of inverse change was 0.4. The results are shown for (A) ECEA, (B) GSEA, and (C) ORA.*

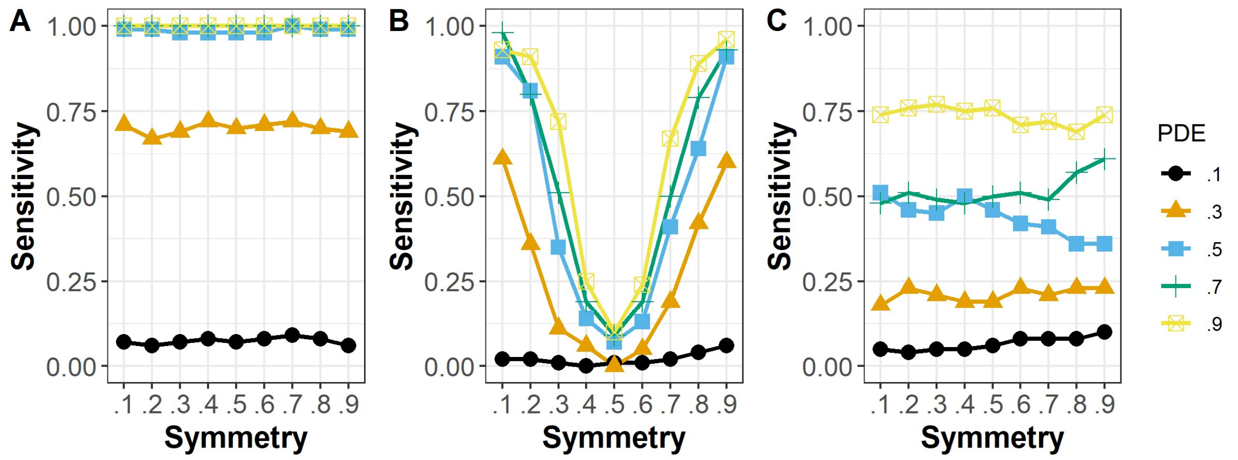


*Figure S64: The proportion of inversely changed pathways detected by each method, when the probability of inverse change was 0.5. The results are shown for (A) ECEA, (B) GSEA, and (C) ORA.*

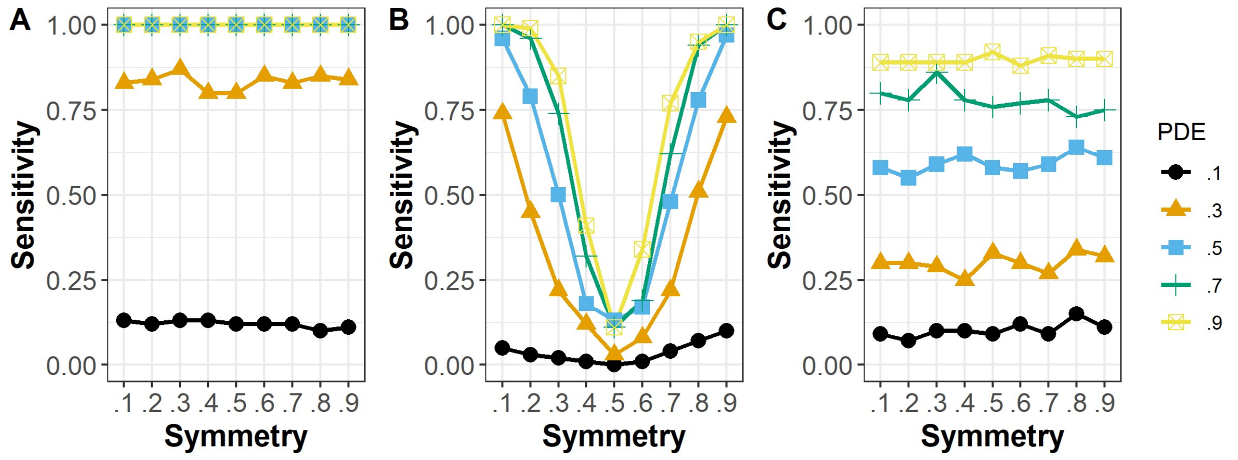

*Figure S75: The proportion of inversely changed pathways detected by each method, when the probability of inverse change was 0.6. The results are shown for (A) ECEA, (B) GSEA, and (C) ORA.*

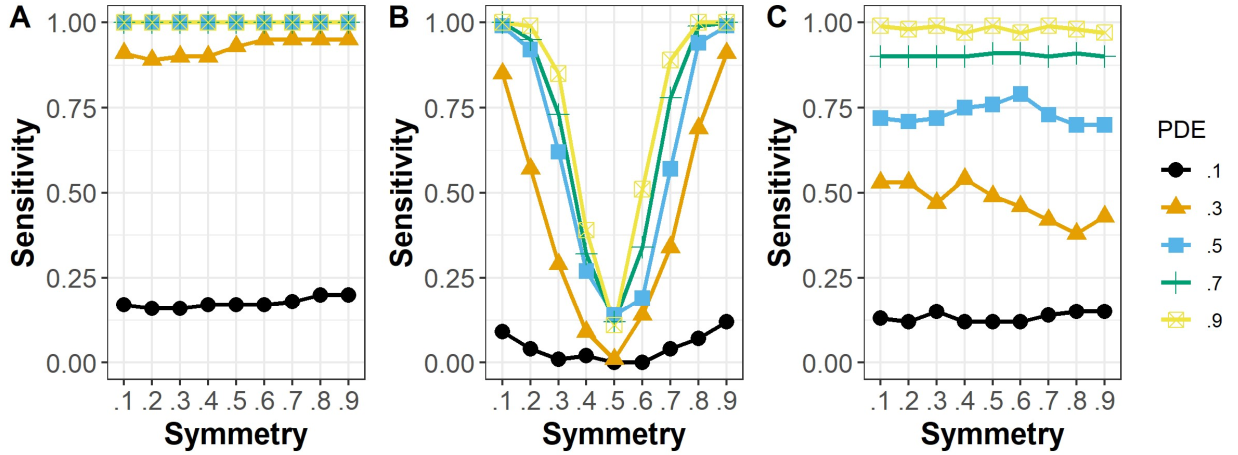


*Figure S86: The proportion of inversely changed pathways detected by each method, when the probability of inverse change was 0.7. The results are shown for (A) ECEA, (B) GSEA, and (C) ORA.*

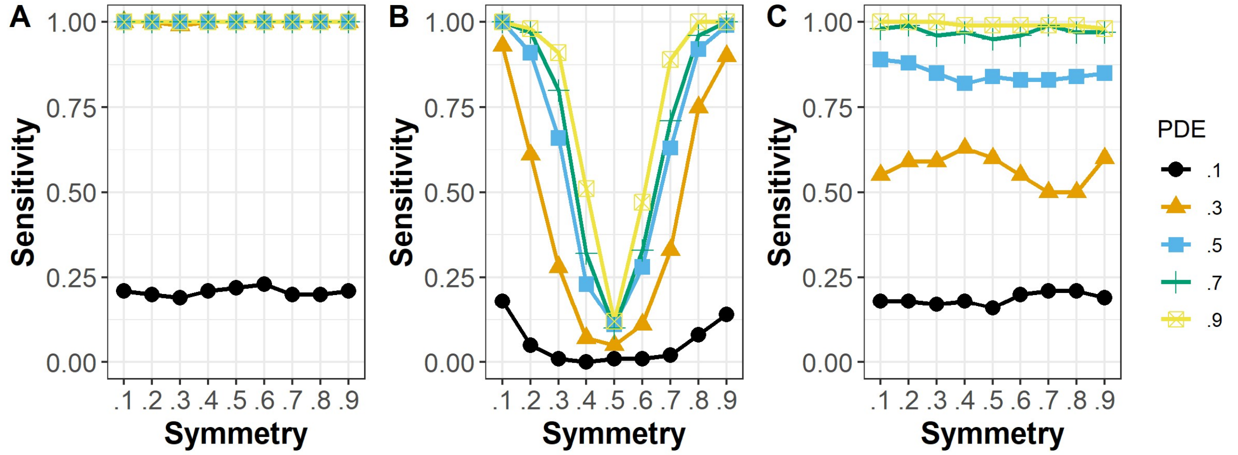


*Figure S97: The proportion of inversely changed pathways detected by each method, when the probability of inverse change was 0.8. The results are shown for (A) ECEA, (B) GSEA, and (C) ORA.*

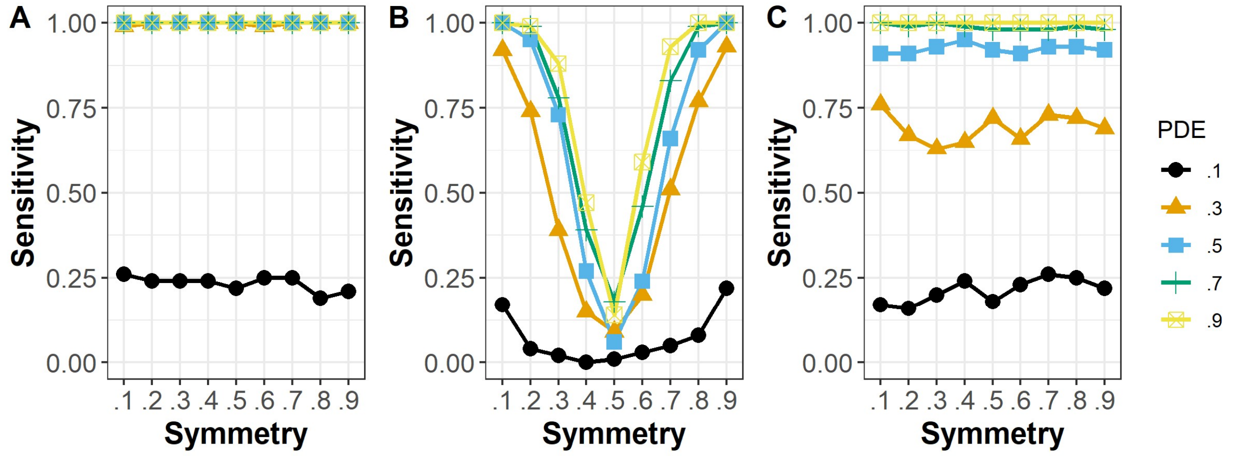

*Figure S108: The proportion of inversely changed pathways detected by each method, when the probability of inverse change was 0.9. The results are shown for (A) ECEA, (B) GSEA, and (C) ORA.*
